# Large-scale computational analyses of gut microbial CAZyme repertoires enabled by Cayman

**DOI:** 10.1101/2024.01.08.574624

**Authors:** Q. R. Ducarmon, N. Karcher, H.L.P. Tytgat, O. Delannoy-Bruno, S. Pekel, F. Springer, C. Schudoma, G. Zeller

## Abstract

Carbohydrate-active enzymes (CAZymes) are crucial for digesting glycans, but bioinformatics tools for CAZyme profiling and interpretation of substrate preferences in microbial community data are lacking. To address this, we developed a CAZyme profiler (Cayman) and a hierarchical substrate annotation scheme. Leveraging these, we genomically survey CAZymes in human gut microbes (n=107,683 genomes), which suggests novel mucin-foraging species. In a subsequent meta-analysis of CAZyme repertoires in Western versus non-Western gut metagenomes (n=4,281) we find that non-Western metagenomes are richer in fibre-degrading CAZymes despite lower overall CAZyme richness. We additionally pinpoint the taxonomic drivers underlying these CAZyme community shifts. A second meta-analysis comparing colorectal cancer patients (CRC) to controls (n=1,998) shows that CRC metagenomes are deprived of dietary fibre-targeting, but enriched in glycosaminoglycan-targeting CAZymes. A genomic analysis of co-localizing CAZyme domains predicts novel substrates for CRC-enriched CAZymes. Cayman is broadly applicable across microbial communities and freely available from https://github.com/zellerlab/cayman.

## Introduction

Carbohydrate-active enzymes (CAZymes) are enzymes that act on glycans and glycoconjugates. They are found throughout the tree of life, but are particularly well-represented in microorganisms and a key factor shaping the metabolic capacity of microbial communities^1^. The human gut microbiota can utilise an enormous diversity of diet-derived (complex) carbohydrates that are otherwise indigestible to the host – as the human genome encodes only 17 catabolic CAZymes^2^. Whilst gut microbes greatly differ in their genomic CAZyme repertoire and preferred substrates, they also express many CAZymes dedicated to degrading host-derived glycans, most importantly mucins lining the intestinal epithelium^2,3^. The *Bacteroides* genus is particularly rich in CAZymes and *Bacteroides* spp. have been shown to flexibly alternate between feeding on mucins and dietary fibre through transcriptional regulation of CAZymes according to their availability in the gut^4^.

Gut microbial carbohydrate metabolism does not only shape microbial communities, but is also crucial for host health. Dietary fibres can be fermented by bacteria into short-chain fatty acids, which promote epithelial barrier integrity and gut health^5,6^. Consequently, fibre-deprived diets have been shown to lead to erosion of the mucus layer as bacteria shift their metabolism to mucus utilisation, which in turn facilitates gut microbes to get closer to the epithelium^3,4^. Compromised barrier integrity is an important feature of gastrointestinal diseases, such as inflammatory bowel disease (IBD) and colorectal cancer (CRC), in which gut microbial CAZymes involved in degradation of host extracellular matrix and mucins are enriched as shown by metagenomic analyses^7,8^, respectively. Given the central role of the gut microbiome as a mediator of dietary effects on host health, understanding how Western lifestyle shapes the CAZyme repertoire can yield important insights into disease processes. Previous work has revealed drastic differences in human gut microbiome composition of Western- and non-Western individuals and provided evidence for the hypothesis that specific microbes and associated functions, including CAZymes, got lost in the process of Westernisation^9,10^. While the binary classification of lifestyles into Western/non-Western may be an oversimplification, the stark contrast has been instrumental to uncover shifts of gut microbiome functions, such as a relative increase of CAZymes facilitating mucin foraging in Western individuals^11^. Despite the crucial roles of CAZymes for both microbial communities and their host, existing studies are limited to few isolate genomes and do not leverage contemporary genomic and metagenomic data resources^2,12,13^. This is the case for comparisons of gut microbial CAZymes between both Western and non-Western populations^11,14^ as well as between diseased and healthy individuals^7,8^.

One underlying reason for the relative scarcity of metagenome-driven CAZyme studies is the lack of scalable and easy-to-use bioinformatics tools: despite the availability of the CAZy database (http://www.cazy.org)^15^ and the annotation framework dbCAN^16^, delineation and quantification of microbial CAZymes in human gut metagenomes is generally performed ad-hoc due to lack of open-source software^8,11,14^. Furthermore, while CAZymes have been divided into the main classes of GH (Glycoside Hydrolase), PL (Polysaccharide Lyase), CE (Carbohydrate Esterases), GT (Glycosyl Transferases), and CBM (Carbohydrate Binding Module), substrate information is more difficult to pinpoint. Substrate information is collected in the CAZy database and through curation efforts of several groups^11,15^, but there can be discrepant classifications and these have to our knowledge not been integrated into computational tools for (meta-)genome annotation.

Here, we developed the first easy-to-use command-line tool to profile CAZymes from shotgun metagenomic data and furthermore provide a manually curated substrate annotation scheme facilitating the interpretation of the resulting CAZyme profiles by grouping CAZyme families into higher-level, biologically meaningful substrate groups. We applied these tools on large-scale bacterial gut genome collections and metagenomic datasets demonstrating their utility for pinpointing bacterial species with specific substrate utilisation patterns (e.g. mucin-foraging) and how glycan substrate utilisation can differ across host lifestyles and health states. Cayman is broadly applicable to (meta-)genomic data from other microbial communities and freely available under https://github.com/zellerlab/cayman.

## Results

### Developing and curating a CAZyme substrate scheme

Our understanding of CAZyme repertoires of microbial communities depends on detailed data about the substrates these enzymes act on. Automated analysis of metagenomic data with respect to CAZymes would thus be greatly facilitated by systematic substrate information. Here we developed such a comprehensive hierarchical substrate annotation scheme for CAZymes (**Methods**) with the aim to overcome limitations of previous efforts that were often incomplete and focussed on glycoside hydrolases (GH) only^11,15^. Here, three authors manually curated available substrate information from http://www.cazy.org/^15^ and related scientific literature for all CAZyme (sub)families of the GH, PL and CBM categories that are represented in dbCAN2 version 9 and formed a consensus on their substrates. This resulted in a table (**Table S1**) which contains, for each individual CAZy (sub)family, several layers of substrate annotation categories that are largely based on previous recommendations for glycan classification^17^. These can be used for interpretation of downstream statistical analysis at various levels of granularity (**Table S2**, **Methods**). For example, starting with CBM75 which binds xyloglucan, our assigned substrate annotations indicate that xyloglucan is a non-starch polysaccharide (NPS) of structural origin that belongs to the hemicellulose group of dietary fibres (**Fig. 1A**).

**Figure 1.**
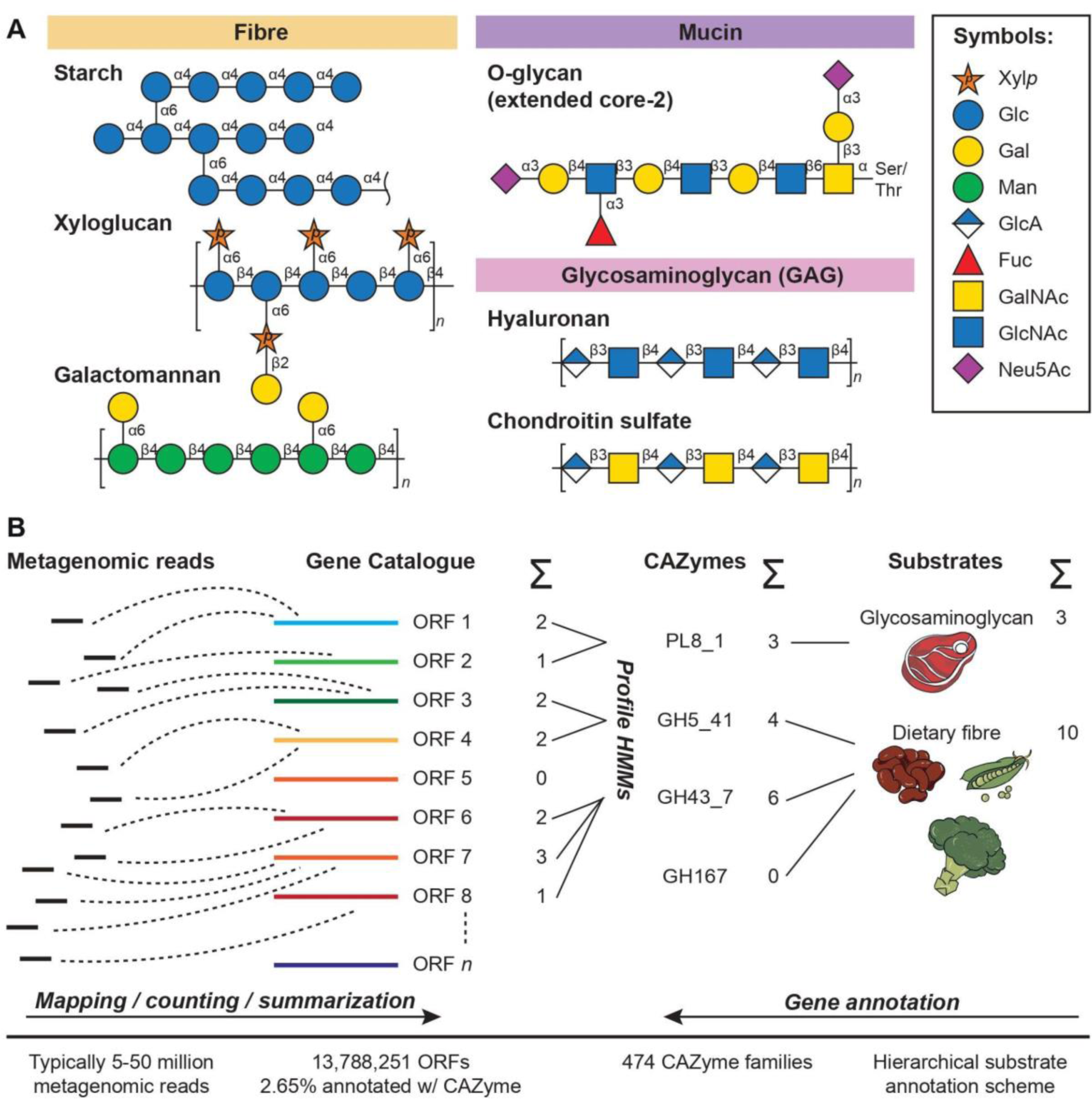
(**A**) Examples of glycan structures and of their classification into high-level substrate categories. Monosaccharide abbreviations: xylopyranose (Xyl*p*), glucose (Glc), galactose (Gal), mannose (Man), glucuronic acid (GlcA), fucose (Fuc), *N*-acetylgalactosamine (GalNAc), *N*-acetylglucosamine (GlcNAc), *N*-acetylneuraminic acid (Neu5Ac).(**B**) Cayman (**C**arbohydrate **a**ctive enz**y**mes profiling of **m**et**a**ge**n**omes) is a computational tool which profiles Carbohydrate-Active Enzymes (CAZymes) from metagenomes via quantification of CAZymes through a gene catalogue. CAZyme abundances are then aggregated/grouped at different levels, facilitated by our curated substrate annotations. CAZyme: Carbohydrate-Active Enzymes, GMGC: Gut Microbial Gene Catalog, ORFs: Open Reading Frames, PL: Polysaccharide Lyase, GH: Glycoside Hydrolase.

### Cayman, a metagenomic CAZyme profiling tool

To be able to routinely quantify CAZymes in metagenomic data, we developed a computational CAZyme profiling tool called Cayman (<u>C</u>arbohydrate <u>a</u>ctive enz<u>y</u>mes profiling of <u>m</u>et<u>a</u>ge<u>n</u>omes). Instead of directly screening for fragments of CAZyme genes in (short) metagenomic sequencing reads, Cayman proceeds by first mapping reads to a gene catalogue, which represents a near-complete complement of microbial genes. Here, for the analysis of human faecal metagenomes, we relied on a recently introduced non-redundant human gut gene catalogue^18^, which we further reduced to ∼13.8 million genes by removing sequences of very low prevalence (0.5% across included metagenomes for gene catalogue construction) and in which we annotated CAZymes using recalibrated profile Hidden Markov models (pHMMs, see **Methods**). This functional profiling via gene-catalogue mapping provides faster and more accurate gene abundance estimates compared to annotating reads themselves^18,19^. Cayman finally calculates length- and sequencing depth-normalised CAZyme family abundances (**Fig. 1B, Methods**) from tallies of mapped reads that can be further summarised using the annotation categories introduced by our curated CAZyme substrate scheme, to e.g. quantify mucin-utilisation CAZymes in a given metagenome and compare across individuals or study populations (**Fig. 1B**).

### Genomic exploration of CAZyme repertoire in human gut microbes

Owing to recent advances in metagenome assembly, genomic resources of the human gut microbiome have substantially grown. To provide an updated view of the CAZyme repertoire of human gut microbes on this basis, we re-annotated genes from 107,683 high-quality Metagenome-Assembled Genomes (MAGs) and isolate genomes^12^. Our analysis confirmed that the genomic potential to metabolically target glycans varies strongly across taxonomy (**Fig. 2A**)^2^. In addition to confirming that key genera from the *Bacteroidetes* phylum, most prominently *Bacteroides* and *Parabacteroides*, have an extensive repertoire of CAZymes, our analysis also highlighted less studied ones, such as *Coprobacter* and *Paraprevotella,* to be particularly rich in CAZymes (213 genes from 80 families and 239 genes from 92 families, respectively). While *Firmicutes* generally showed lower CAZyme diversity (on average 77.2, sd 38.7, CAZymes/genome compared to an average of 181, sd 105, for *Bacteroides*), several specific genera from this phylum exhibited exceptionally rich CAZyme repertoires. Among these, *Hungatella* and *Eisenbergiella* stood out as distinctly rich in CAZymes (with 224 genes from 78 families and 404 genes from 106 families, respectively), but to our knowledge, neither genus has been experimentally studied for their glycan utilisation capabilities (**Extended Data Fig. 1**).

**Figure 2.**
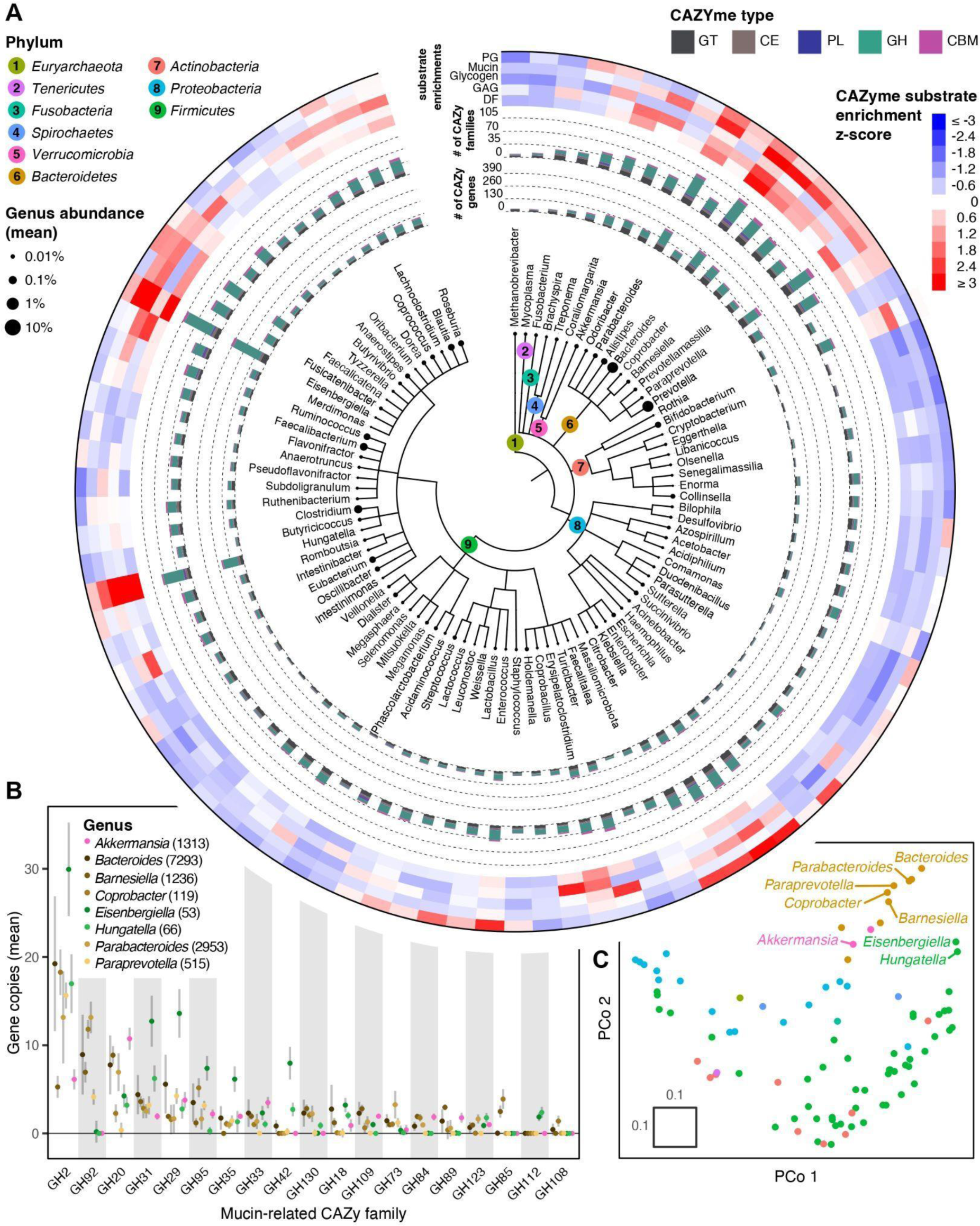
(**A**). Taxonomic tree of 90 prevalent human gut microbial genera (based on n=75,131 genomes) showing their mean relative abundance (leaf tips), number of encoded CAZyme genes (inner coloured bar plot), number of unique CAZyme families encoded (outer coloured bar plot) by type as well as CAZyme substrate enrichments (outermost heatmap). Substrate enrichment was calculated as z-scores of the genus-wise mean total copy number of CAZymes annotated with a specific substrate class. (**B**) Whisker plot of genomic copy numbers of mucin-related CAZymes in genomes of the 8 genera with the highest average mucin-related CAZyme copy number; dot corresponds to mean and bar corresponds to standard deviation and numbers of genomes are given in brackets. (**C**) Ordination plot depicting genomic CAZyme similarity between genera (colour code as in (A)). CAZyme similarity between pairs of genera was calculated as the Jaccard index of genomic absence/presence of CAZymes (**Methods**).

To compare the overall CAZyme repertoire of gut microbial genera, we further performed an ordination based on the pairwise similarity of encoded CAZy families (**Methods**). This shows a trend of genera from the same phylum to cluster based on their overall CAZyme repertoire. Contrary to that, the ordination also grouped *Hungatella, Eisenbergiella* and the CAZy-rich *Bacteroides* together and farthest apart from the gut genera with the fewest CAZyme genes, such as *Actinobacteria* or *Proteobacteria* (**Fig. 2A, C**).

To investigate CAZymes substrate preferences among core gut bacterial genera, we leveraged our substrate annotations to compute enrichment scores of 5 major substrate classes: peptidoglycan (PG), mucin, glycogen, glycosaminoglycan (GAG) and dietary fibre (DF) (**Methods**, **Fig. 2A**). This analysis confirmed previously well-described mucin-foragers such as *Akkermansia, Alistipes* and *Bacteroides* to possess many different mucin-targeting CAZyme genes. The relatively poorly characterised *Barnesiellaceae* family included two genera, *Barnesiella* (represented in the human gut by the singular species *B. intestinihominis*) and *Coprobacter* (represented in the human gut by the species *C. fastidiosus and C. secundus*) that were also strongly enriched in mucin-targeting CAZymes. While *B. intestinihominis* has recently been described as a mucin specialist that can grow exclusively on mucin O-type glycans^20^, this has to our knowledge not been reported for *Coprobacter* spp., which our results suggest for the first time to be a prolific mucin forager. Lastly, we found *Hungatella* and *Eisenbergiella* genomes to be strongly enriched in mucin-targeting CAZymes (**Fig. 2A**, **Fig. 2B**) with an overall CAZyme repertoire similar to known mucin foragers (**Fig. 2C**). In particular *Eisenbergiella* exhibited the highest number of genomic copies for several mucin-targeting GHs, including α-fucosidases (GH29 and GH95), highlighting the promise of further characterization of its mucin-foraging capabilities. While experimental data on mucin utilisation for *Eisenbergiella tayi* – the only *Eisenbergiella* species found in the human gut – is lacking, it has been observed to increase after supplementing mucin to culture media and bioreactors^21,22^. Taken together, our analysis gives a systematic overview of carbohydrate-related metabolic potential of human gut bacteria and showcases the utility of our method for genomic analysis of microbial CAZyme repertoires and substrate preferences.

#### Meta-analysis of Western versus non-Western gut metagenomes

To investigate if there are systematic CAZyme repertoire differences across human populations, we applied Cayman to publicly available Western and non-Western gut metagenomes (n=3,189 from Western and n=1,092 from non-Western subjects). These metagenomes are derived from populations of geographically diverse locations, with the Western metagenomes coming from North-American, European and Asian individuals and the non-Western metagenomes from Central-American, South-American, African, Asian and Oceanian individuals (**Table S3**). Principal Coordinate Analysis of CAZy profiles shows clear separation between Western and non-Western microbiomes (PERMANOVA, R^2^ = 0.073, p = 0.001), highlighting pronounced abundance differences of CAZyme families between these groups (**Fig. 3A**).

**Figure 3.**
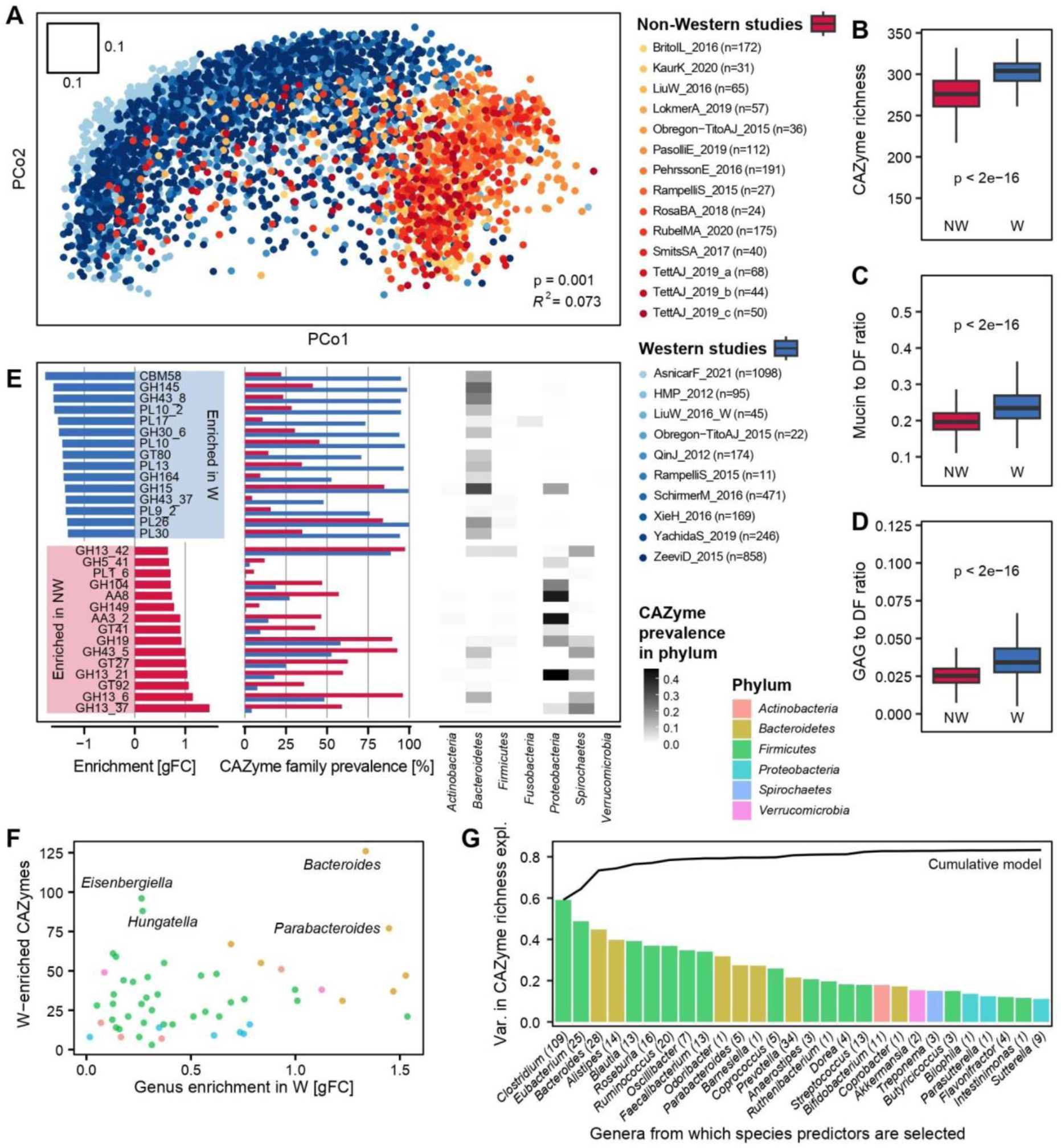
(**A**) Principal Coordinates Analysis using pairwise Canberra distances between gut microbiome CAZy profiles of Western (W) and non-Western (NW) individuals (n=4,281, see key for studies, some of which contain both Western and non-Western metagenomes, and **Extended Data Fig. 2** for ordinations based on other dissimilarity measures). PERMANOVA was applied for statistical testing of group differences. (**B**) Boxplots of the number of unique CAZymes between Western (n=3,189) and non-Western (n=1,092) individuals. (**C**) Boxplots of the ratio between the total abundance of mucin-targeting and dietary fibre-targeting CAZymes in Western- and non-Western microbiomes. (**D**) Boxplots of the ratio between the total abundance of GAG-targeting and dietary fibre-targeting CAZymes in Western- and non-Western microbiomes. The boxplot centre value corresponds to the median, the box indicates the interquartile range, and whiskers extend to 1.5 times the interquartile range, while dots indicate outliers. Significance was tested using unpaired, two-sided Wilcoxon tests. (**E**) Bar plots showing the 15 most strongly Western-enriched (blue) and 15 most strongly non-Western enriched (red) CAZyme families based on generalised fold change (gFC), their prevalence and a heatmap of mean phylum-level prevalences of these families based on genomic data. (**F**) Genus enrichment in Western individuals plotted against the number of Western-enriched CAZymes encoded per genus (calculated as explained in E). (**G**) Variance in community CAZyme richness explained by microbial species abundances grouped by genus. Barplots show the R^2^-values of linear models based on mOTU^23^ abundances from within a single genus. Line plot indicates the R^2^-value of a cumulative model based on all mOTUs up to a given genus. The numbers in parentheses correspond to the number of mOTUs added for the given genus (**Methods**).

It has previously been described in smaller-scale studies that non-Western microbiomes generally have a higher CAZyme diversity compared to Western ones, a phenomenon mostly attributed to the substantial reduction of dietary fibre content in Western diets^9–11^. In sharp contrast to the expectation formed on these reports, we found a consistently higher number of unique CAZymes in Western compared to non-Western populations, even when only considering CAZymes involved in dietary fibre metabolism (**Fig. 3B, Extended Data Fig. 3, Extended Data Fig. 4**). While we noticed a significantly higher read mapping rate in Western compared to non-Western metagenomes (**Extended Data Fig. 5**), an additional comparison of Western and non-Western samples with similar read mapping rates rendered this explanation unlikely (**Extended Data Fig. 6**).

To investigate to which extent the CAZyme family abundances reflect dietary differences, we leveraged our substrate annotation scheme to compare Western to non-Western CAZyme repertoires. Generally, Western diets are characterised by high amounts of animal-based products and low fibre content, while many non-Western diets are instead characterised by a high intake of fibre-rich, plant-based foods, although large variation exists within non-Western diets^24–26^. Our substrate ratio analyses confirmed previous findings that the abundance ratio between mucin-targeting and dietary fibre-targeting CAZymes is higher in Western compared to non-Western metagenomes (**Fig. 3C**). Similarly, Western metagenomes have a higher abundance ratio of CAZymes targeting GAGs (animal-derived glycans) relative to those targeting dietary fibre (**Fig. 3D**). Combined, these findings confirm an increased richness and abundance of gut microbial metabolic potential for the degradation of host and animal-derived glycans in Western metagenomes.

To resolve differences in individual CAZyme families between Western and non-Western subjects, we subjected each family to differential abundance testing using linear models (**Fig. 3E**). This analysis indicated a large number of CAZymes to differ significantly in abundance (adjusted p < 0.05 for 397 / 457 CAZyme families) and in prevalence (**Fig. 3E**). Among the most non-Western enriched CAZymes, we observed four families (GH13_37, GH13_6, GH13_21, GH13_42) that target resistant starch, which could be explained by the generally higher intake of whole grains and legumes in non-Western populations^27^. To attribute the differentially abundant families to their likely taxonomic origin, we compared the CAZyme prevalence over different phyla (**Methods**). This revealed that many CAZymes enriched in non-Western individuals are prevalent in *Proteobacteria* and *Spirochaetes* (**Fig. 3E**). The latter phylum has previously been labelled a ‘vanish’ taxon, meaning that it is common in non-Westernised gut microbiomes, but is lost or strongly reduced in more Westernised individuals^9^. On the other hand, the most Western-enriched CAZyme, CBM58, is part of the SusG protein, which is crucial for starch utilisation in *Bacteroides* species. This is consistent with the taxonomic enrichment of *Bacteroides* in Western individuals (**Extended Data Fig. 7**).

To assess whether *Bacteroides* expansion could fully explain the increased CAZyme richness in Western populations, we first asked how many Western-enriched CAZymes each (Western-enriched) bacterial genus could contribute to the community repertoire. This analysis indeed singled out *Bacteroides* to have by far the largest number of Western-enriched CAZymes (n=126, **Fig. 3F**). We next investigated to which extent the genomic CAZyme repertoire of individual taxa or a small number of key taxa would be predictive of the whole community’s CAZyme richness across human populations. To this end, we constructed linear regression models for CAZyme richness from genus abundances with stepwise increasing sets of taxonomic predictors (forward variable selection, see **Methods**) (**Fig. 3G**). In this analysis CAZyme richness could be predicted well by a few genera (R2-value of 0.76 with top 5 predictive genera). Interestingly, while *Bacteroides* was among these predictors, *Eubacterium* and *Clostridium* were even more predictive of community CAZyme richness. Taken together, these results indicate that while Western-enriched *Bacteroides* is genomically the richest in Western-associated CAZymes and thus likely responsible for much of the increased CAZyme richness observed in Western individuals, its abundance alone is not sufficient to explain community CAZyme content. Instead, there appear to be several key taxa that together shape the community CAZyme pool; their taxonomic variation across individuals indeed explains most of the inter-individual differences in CAZyme richness across human populations.

To connect taxonomic composition to community CAZyme repertoire more broadly and at higher resolution, we built linear regression models to predict the abundance of each CAZyme family from species abundance profiles within each gut microbial genus. We trained these models separately for Western and non-Western populations to be able to compare the associations (**Methods**). We found strong pairwise associations between taxonomic and CAZyme abundances and generally, as expected, we observed the contributions to CAZyme abundances of highly abundant genera, such as *Bacteroides* and *Prevotella*, to outweigh those of rare, low-abundant genera (**Fig. 4A**). However, we noted exceptions to this trend, for example *Bifidobacterium* and GH13_3 (**Fig. 4B**). While this CAZyme family has not been experimentally characterised in any gut microbe yet^15^, our analysis shows GH13_3 abundance to be very well predictable from *Bifidobacterium* spp. abundances, suggesting this genus to be the sole contributor to the GH13_3 family in both Western and non-Western human gut microbiomes. Secondly, we observed that some associations strongly differ between Western and non-Western individuals despite taxa being similarly abundant: For example, *Collinsella* is strongly predictive of CAZyme GH13_30 in non-Western individuals but markedly less so in Western individuals (**Fig. 4C**). The fact that *Collinsella* does not differ strongly in abundance suggests that other taxa contribute to this CAZyme pool in Western but not non-Western individuals. A similar but ecologically distinct example is GH95 (encoding a α-fucosidase, needed for mucin foraging): While in Western individuals this CAZyme family has a strong association with *Bacteroides* (**Fig. 4D**), but not with *Prevotella* (**Fig. 4E**), the opposite pattern was observed for non-Western individuals. This suggests that in the process of Westernisation *Bacteroides* as an expanding taxon has taken over functionalities originally provided by *Prevotella*. This example is especially interesting given that *Prevotella* is generally not regarded as a mucin forager. Additionally, no enzyme from *Prevotella* has been experimentally identified as GH95, even though this family has been characterised in numerous gut microbes (including *Akkermansia*, *Bacteroides*, *Roseburia* and *Ruminococcus*)^15^. As a final example, *Akkermansia* is highly predictive of GT31 (a family involved in synthesis of many glycoconjugates) in Western but not in non-Western individuals (**Fig. 4F**), showing that *Akkermansia* is the principal carrier of this CAZyme family in Western individuals but not in non-Western individuals.

**Figure 4.**
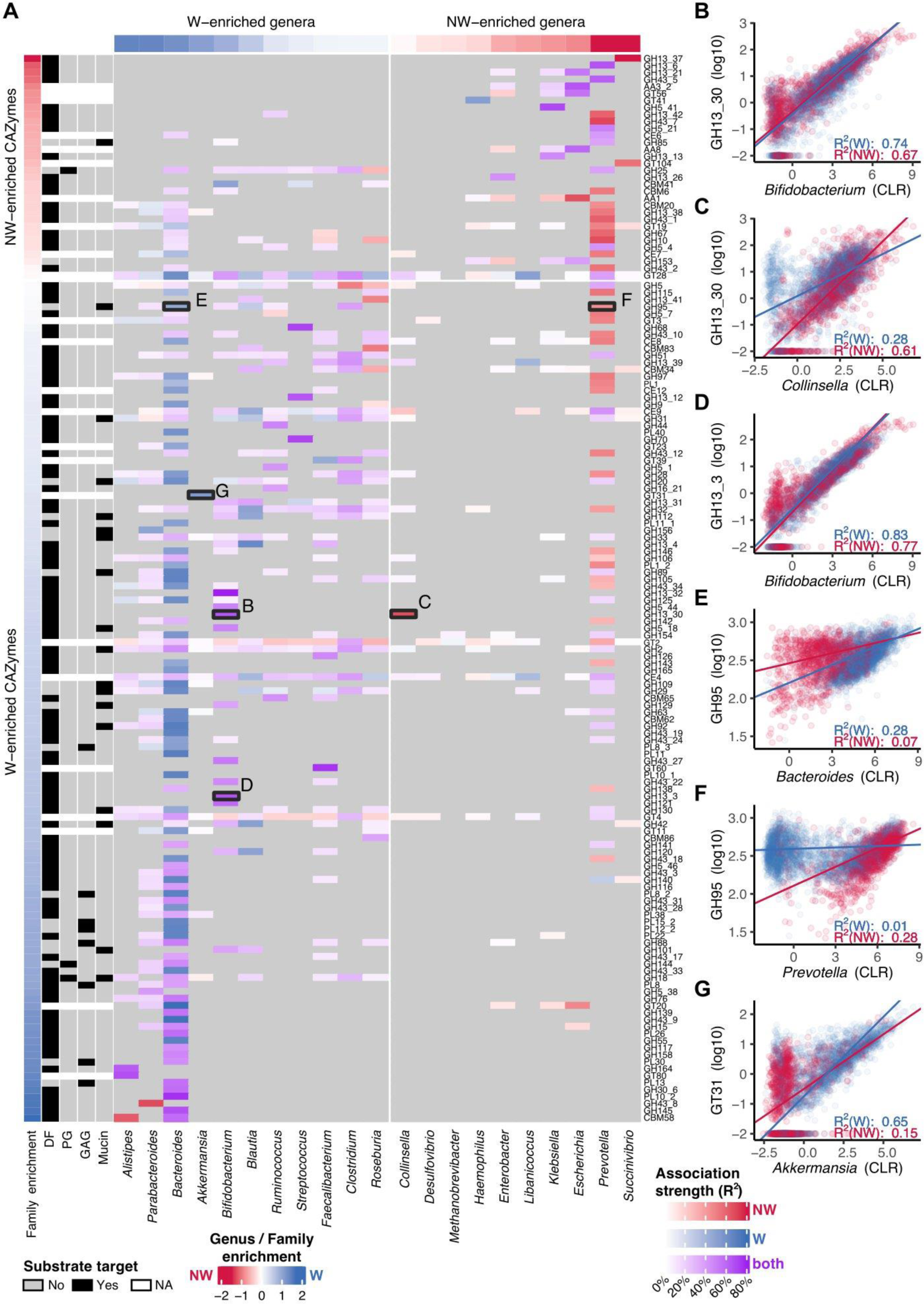
(**A**) Heatmap showing predicted taxonomic contributions to CAZyme abundances in Western-(n=3,189) and non-Western (n=1,092) gut microbial communities. Coloured cells depict the R^2^-value of multivariable linear models in which log10-scaled CAZyme coverage values were predicted from all CLR-transformed species (mOTU) abundances within a given genus. Blue and red cells indicate better fits in Western or non-Western models, respectively, while purple cells indicate similar fits for Western- and non-Western models (R^2^-values within 2-fold difference). Grey cells indicate models with an insignificant fit after multiple testing correction at 1% FDR. Western vs non-Western enrichments for taxa and CAZy families were defined as previously (see Fig. 3) and substrate annotations for CAZymes are included as leftmost columns. (**B-G**) Genus-level scatter plots between taxonomic abundances and CAZyme abundances; in all of these cases genus- and mOTU-level associations with CAZyme abundance were similar (**Extended Data Fig. 8**), linear fits are indicated by R^2^-values (**Methods**).

Taken together, our results show the major taxonomic contributors to community CAZyme pools. By contrasting these contributions between Western and non-Western individuals, we reveal novel facets of how Westernisation has shaped the gut microbiome at both the taxonomic and functional layer and how these are interconnected.

#### Meta-analysis of colorectal cancer case-control studies

A Western lifestyle is a major driver of many common diseases, including colorectal cancer (CRC). For this condition, differences in gut microbial composition between cancer patients and controls are well understood^28^ and differences in their CAZyme repertoire have also been reported in individual studies^8^. Here we sought to provide a detailed assessment of gut CAZyme repertoires through the application of Cayman in a meta-analysis of CRC case-control studies including 1,998 metagenomes in total (n=968 CRC patients and n=1,030 controls, **Table S4**) from 4 continents and 10 different countries.

We first computed differentially abundant CAZyme families per study (**Fig. 5A**, **Methods**). This analysis revealed broadly consistent patterns of CAZyme enrichments and depletions in CRC patients across studies (**Fig. 5A**), but in line with a previous meta-analysis, several datasets did not show significant differences between CRC and control for various reasons^28^.

**Figure 5.**
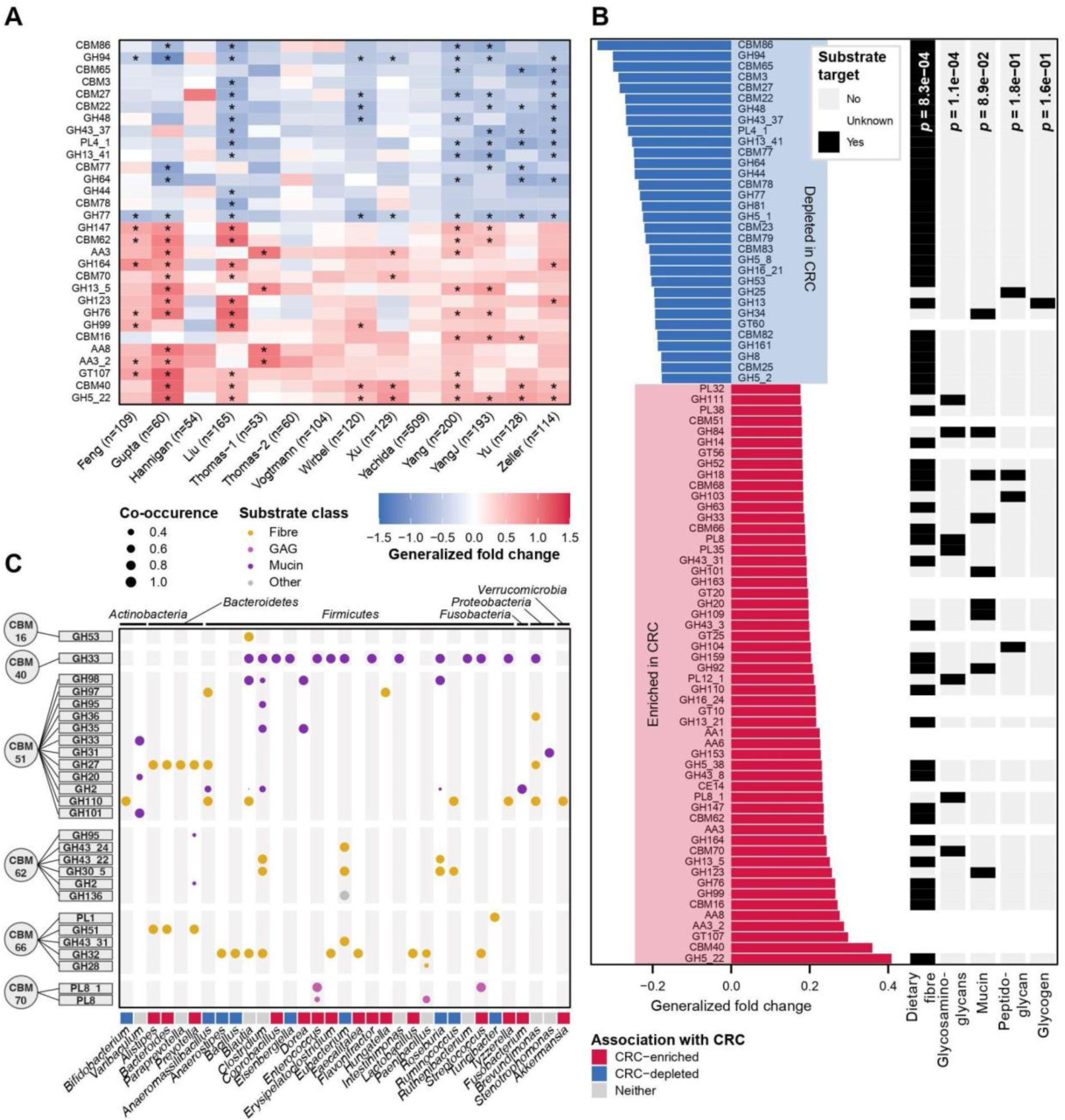
(**A**) Heatmap with univariate associations per CRC study. Asterisk signifies per-study FDR-corrected significance (BH-adjusted p-value < 0.05). Top and bottom 15 families were selected based on the order of CAZyme families from the meta-analysis in panel B (all having BH-adjusted p-value < 0.05). (**B**) Meta-analysis of CAZyme enrichments over all studies using LMMs with study as a random effect. Subsequently, all CAZyme families were subjected to gene-set enrichment analysis (GSEA) to identify overrepresented substrate groups. (**C**) Heatmap depicting CRC-enriched CBMs co-occurring with catabolic CAZyme families within the same gene. Co-occurrences were calculated using a modified jaccard on bacterial genomic data (**Methods**). Dot size corresponds to the strength of association and colour indicates substrate annotation of the enzymatic domain.

To perform a meta-analysis of differentially abundant CAZyme families while accounting for study heterogeneity, we fitted linear mixed models predicting cancer state from CAZyme profiles with study as a random intercept. Out of 459 CAZymes included, 282 were significant at 5% FDR (**Fig. 5B**). We next aimed to identify potential community-wide shifts in carbohydrate substrate preferences by leveraging our substrate scheme in a gene-set enrichment analysis (**Methods**). This analysis showed that CRC metagenomes were depleted in CAZyme families involved in dietary fibre metabolism (p = 1.1e-4), while they were strongly enriched in CAZymes involved in GAG metabolism (**Fig. 5B** and p = 8.3e-4) with a trend toward enrichment for mucin metabolism (p = 8.9e-2). Mucin to DF and GAG to DF ratios further confirmed this finding, with both being increased in CRC patients (**Extended Data Fig. 9**). Taken together, these results showed a clear enrichment of CAZymes involved in host- and animal glycan metabolism in CRC metagenomes, while CAZymes involved in dietary fibre metabolism were depleted. These findings are in line with earlier reports^8,28^ and dietary epidemiological risk factors for CRC development^29–31^ and might in part be a consequence of shifts in dietary patterns as well as ecological adaptations of the gut microbiome to CRC.

Interestingly, for some of the most strongly CRC-enriched CAZymes, our substrate curation efforts (see above and **Methods**) could not pinpoint a substrate. Given that CBMs tend to co-occur with functionally linked enzymatic domains (GHs or PLs) on the same gene^32^, we investigated if such co-occurrence patterns could yield hints about their substrate binding functions. To this end we determined co-occurrence between CRC-enriched CBMs and enzymatic CAZyme families in our bacterial genome collection (**Methods**) and identified consistently co-occurring pairs (**Fig. 5C**). Some of the co-occurrence patterns we found were taxonomically conserved across many bacterial phyla (e.g. CBM40-GH33 and CBM51-GH110), while others are more restricted (e.g. CBM16-GH53 being specific to *Blautia* and CBM51-GH101 being specific to *Varibaculum*). Furthermore, some CBMs, e.g. CBM51, co-occurred with a variety of different catabolic CAZyme families, while others, including CBM12 and CBM40, co-occurred with one or few specific enzymatic families.

Leveraging the genomic co-occurrence data to deduce function in the ‘guilt-by-association’ paradigm, we next compared the substrate annotations of the CBM and GH/PL domains co-occurring on the same genes. This analysis confirmed several previously reported associations: For example, the consistent and specific co-occurrence of CBM40 and GH33 across more than a dozen gut bacterial genera indicated that CBM40, and not only GH33, facilitate mucin foraging^33,34^. Similarly, CBM70 specifically co-occurred with PL8 (and its subfamily PL8_1), a known GAG-degrading enzyme, consistent with a demonstration of CBM70 binding the GAG hyaluronan^35^. While our curated substrate annotations showed CBM51 to have blood group antigens as the main substrate, we found it co-occurring with a large number of CAZymes previously implicated in mucin utilisation. Indeed, an additional literature search confirmed the mucin-binding capability of CBM51^36^. Finally, from our observation that all GH families involved in fibre metabolism that co-occur with CBM51 have α-galactose among their substrates, we predict that CBM51 also binds this glycan.

Taken together, our genomic co-occurrence analysis combined with our substrate annotations revealed (i) large variation in conservation of CAZyme domain pairs across gut microbes, (ii) co-occurring CBM-GHs often share similar substrate specificities and (iii) that substrate specificity of unannotated CAZymes can be predicted using a ‘guilt-by-association’ approach. Taking this strategy, we uncovered novel associations that could hint at previously overlooked mucin-utilisation functions in human gut microbes.

## Discussion

In this work we present Cayman, the first open-source software tool for profiling CAZymes from metagenomic data, along with a hierarchical substrate annotation for CAZymes. Cayman facilitates functional profiling to complement the routine taxonomic analysis of shotgun metagenomes. The efficient profiling approach via metagenomic gene catalogues enabled the application to (meta)genomic data sets of much larger scale and is in contrast to most reported CAZymes profiling efforts that are typically more computationally demanding (as usually each sequencing read or assembled contig is annotated). One important previous study characterised the CAZyme repertoire of the gut microbiome on the basis of an artificial ‘mini-microbiome’ that was constructed from 177 isolate genomes, which were subsequently annotated with CAZymes information^2^. This approach comes with crucial limitations for generalising the findings to *in vivo* microbiomes: First, several of the included genomes were of enteropathogens not normally seen in a gut microbiome^2^. Second, this setting did not reflect realistic abundances of gut bacteria^2^. As an important limitation that our analysis shares with this and previous efforts, CAZyme and substrate annotations are transferred via sequence homology, as experimental data linking the CAZyme to their substrate is lacking for the vast majority of (gut) microbes.

Nevertheless, Cayman enables large-scale surveys of the gut microbial CAZyme repertoire across a large number of human gut metagenomes to reveal how it is shaped by lifestyle and disease. The comparison between Western and non-Western individuals and between cancer patients and tumour-free individuals revealed interesting similarities. Both Western lifestyle and CRC, that is epidemiologically associated with a Western diet (rich in meat and low in fibre), showed a reduction of CAZymes targeting fibre as substrates. As such, these gene families could potentially serve as a gut microbial marker for compromised gut health.

To infer how individual microbial taxa contributed to the community CAZyme gene pool, we used linear models, inspired by previous work on modelling microbe-metabolite relationships^37^. Our results from comparing Western to non-Western metagenomes suggest that highly abundant *Bacteroidetes* genera *(Bacteroides* in Western individuals and *Prevotella* in non-Western individuals) make key contributions to the community CAZyme pool, and that additionally specific CAZymes are contributed by genera that are less prevalent and more variable across individuals and human populations. Altogether, this strategy thus appears suitable for pinpointing the taxonomic contributions to specific CAZyme abundances in microbial communities.

Our comparison between Western and non-Western gut microbiomes across a large number of geographically diverse study populations revealed the unexpected observation of higher CAZyme richness in Western individuals, which seemingly contradicts current consensus^9,11,14^. There are multiple possible explanations for this discrepancy that support our conclusion: First, we used a larger and more diverse collection of both Western and non-Western datasets than previous studies, in order to obtain more generalisable results. Previous work undersampled Western populations in particular: for instance, Smits et al. compared exclusively against the data from the first phase of the Human Microbiome Project^38^, which in our re-analysis shows an unusually low number of unique CAZymes consistent with a previously observed very low taxonomic diversity^39^. Second, utilisation of a gene catalogue in Cayman as compared to direct CAZyme annotation of reads^11^ or assembled contigs^14^ is expected to yield more accurate results. This is on the one hand because genes (and not just short reads) are annotated in a comprehensive catalogue^18^, and on the other hand because short reads are mapped competitively against the whole catalogue (not only CAZyme-annotated genes), which prevents spurious CAZyme assignments in cases where a read better matches a similar – but functionally distinct – gene sequence without a CAZyme annotation. Lastly, previous studies have not directly assessed CAZyme richness, but rather their (Shannon) diversity, which is also influenced by the evenness of their abundances^9,11^ or gene richness (i.e. the number of ORFs annotated with a CAZyme)^14^. Biologically, increased CAZyme richness in Western individuals is not entirely implausible given the year-round availability of a wide variety of foods (containing diverse types of fibres) in contrast to the availability of specific foods in traditional populations being much more restricted by seasonality and geographic proximity^11,40^. To better rationalise the functional differences between Western and non-Western microbiomes, we will have to obtain more comprehensive data on how dietary ingredients promote the growth of specific bacteria and more systematic experimental data on glycan utilisation capabilities of microbial genes and species.

In conclusion, we present the first easy-to-use command-line tool for profiling CAZymes from metagenomes. We demonstrate how it facilitated the discovery of several novel aspects of CAZyme biology of the human gut microbiome leveraging large-scale metagenomic data.

We anticipate that Cayman will be broadly useful for the study of diverse microbial communities and their glycan metabolism.

## Methods

### Generation of CAZyme (sub)family module sequence set

In order to obtain family-wise CAZy modules, we first downloaded CAZy sequences (http://bcb.unl.edu/dbCAN2/download/Databases/V9/ dbCAN HMMdb release 9.0 and CAZyDB released on 07/30/2020) from a total of 676 families and subfamilies. We then extracted CAZy modules from these sequences using the dbCAN pHMMs (from the same dbCAN2 release) using an E-value threshold of 1e-15 and a coverage threshold of 0.35 (default cut-offs on dbCAN server). Next, we generated family-wise multiple sequence alignments of module sequences using mafft-linsi^41^ on representative module sequences obtained from mmseqs2 with --easy-cluster --min-seq-id 0.99 --cov-mode 0 -c 0.5^42^. Default parameters were used unless stated otherwise.

### Optimization of pHMM p-value cutoffs

We set out to determine family-specific sequence similarity cutoffs for pHMMs for optimised detection of CAZymes in large sequencing datasets. To this end, we constructed and evaluated pHMMs for each CAZy family individually using blocked cross validation: For each CAZy family and cross-validation fold, we divided family sequences into training (∼80%) and testing sets (∼20%): The training set was used to build the family pHMM and the test set was used as positive instances at testing time, while sequences from other CAZyme modules were used as negative instances (more details see below). When building pHMMs from training sequences we omitted sequences shorter than 80% of the median module length. For testing we used entire gene sequences (instead of family module sequences for training pHMMs) since they represent a more realistic search space. Training and test fold generation was done in a blocked fashion, where similar sequences remain together in either training or test set. This was done to minimise information leakage from the training set into the test set. To define blocking groups, module sequences within each family were clustered at 60% sequence identity using mmseqs2 (arguments easy-cluster, --min-seq-id 0.60, --cov-mode 0, -c 0.5). Folds were designed in such a way that test sets never overlap with each other.

We evaluated families differently depending on their hierarchy of subfamilies: When evaluating a family without subfamilies, we considered corresponding family sequences as positive instances and all other sequences as negative instances. When evaluating a family with subfamilies, we proceeded as above but additionally considered all subfamily sequences under that family as positive instances. When evaluating subfamilies, we ignored upstream family sequences and considered all other families (including sister subfamilies) as negative instances. In all cases we furthermore utilised non-CAZy sequences from UniProt as additional negative instances. These sequences were obtained in the following manner. First, all manually curated sequences (Swiss-Prot) with an annotation score of five out of five (indicating experimental evidence at protein level) were downloaded (n=54,978 sequences, March 5th 2021). We subsequently filtered out all sequences with an annotated Enzyme Commission (EC) number present in CAZy, yielding 51,507 sequences. Since CBMs are non-catalytic (and thus have no EC number), we did not add UniProt sequences as negative instances when we evaluated pHMMs for CBMs.

This setup was applied to the 527 CAZyme families that had at least 50 sequences in the CAZy database. For the 130 families which had less than 50 sequences, but 5 or more sequences or where less than 5 blocking groups existed, cross validation was performed without blocking. 11 families had less than 5 sequences and were not cross-validated in this manner but instead were used with the median optimised p-value cutoff of the corresponding CAZyme class. This way we derived cutoffs for a total of 668 CAZy families.

Finally, we determined family- and fold-wise optimal p-values by iterating over p-value thresholds and choosing the p-value that maximises the F1-Score. The F1 score was calculated using the follow formula: 2*(recall*precision) / (recall + precision). Recall was calculated using the formula: TP / (TP + FN) and precision using TP / (TP + FP). We optimised p-value cut-offs in this manner for 647 CAZy families and the 21 CAZy families with an F1 score of <0.5 were removed, as these CAZyme families could not be reliably detected using HMMs.

### Annotating novel sequences

In order to annotate new genomic sequences, we used PyHMMER^43^ to run all rebuilt pHMMs against the new genomic sequence set and filtered the hits using the family-wise optimised p-value cutoffs (see above). Next, we kept only those residues where at least half the fold-specific HMMs (rounded up) yielded a significant hit. Finally, we merged overlapping regions to obtain the annotations as simple coordinates on a genomic sequence.

### Substrate annotation scheme design

To design a meaningful, hierarchical substrate scheme, we aimed to capture for an extensive list of glycans, the origin of the glycan, its function in the organism from which it originated (storage or structural glycan) and its function at destination (especially relevant for mammalian gut environments), which was further subdivided into 3 different categories. The 3 subcategories of function at destination were based on recommendations^17^ that divide dietary fibre into several subclasses (e.g. whether a glycan derives from a non-starch polysaccharide (NSP) or a resistant starch and in which category the class of poly- or oligosaccharide falls). We then applied the same logic to glycans that are not dietary fibres, like glycosaminoglycans and mucin-associated glycans. After having constructed a complete table for selected glycans (**Table S2**) with a hierarchical substrate design, we continued to manually curate all CAZyme (sub)families in the CAZy database. This annotation effort led to glycan-specific annotations of each individual CAZyme (sub)family. By subsequently mapping our **Table S2** onto all individual CAZy (sub)families, we then obtained a large table (**Table S1**) that, for every individual CAZyme family, contained information on the hierarchical categories we designed in **Table S2**. All substrate annotations were initially performed by Q.D. and H.T. and were subsequently, independently validated by O.D-B.

### Genomic annotation of human gut microbial CAZymes

Genomic analysis was on based all high-quality genomes (Completeness > 90%, Contamination < 5%) from Almeida et al.^12^, amounting to a total of 6,456 isolate genomes and 101,229 Metagenome-Assembled Genomes (MAGs) which we annotated with our pipeline. The taxonomic tree in Figure 2 shows 90 prevalent human gut bacterial genera, defined as follows: Based on Westernized and non-Westernized control samples (See section “Western and non-Western datasets”, **Table S4**), we first computed mOTUs that are more than 5% prevalent and whose maximum relative abundance exceeds 1% in at least one sample. We then computed a taxonomy based on all genera that contain at least one of those mOTUs. We defined a CAZyme family to be present in a genus if at least 20% of genomes within that genus carried at least one copy of the family.

### Pre-processing of metagenomic datasets

Prior to functional profiling, raw reads were subjected to the following protocol with bbduk (bbmap-version 38.93): 1) low quality trimming on either side (qtrim = rl, trimq = 3), 2) discarding of low-quality reads (maq = 25), 3) adapter removal (ktrim = r, k = 23, mink = 11, hdist = 1, tpe = true, tbo = true; against the bbduk default adapter library) and 4) length filtering (ml = 45). The cleaned reads were screened for host contamination using kraken2 (version 2.1.2)^44^ against the human hg38 reference genome with ribosomal sequences masked (SILVA 138).

### Cayman profiling steps for obtaining CAZyme abundances from cleaned metagenomic shotgun sequences

As input to Cayman, cleaned metagenomic shotgun reads are mapped to the habitat-specific Global Microbial Gene Catalogue^18^ (GMGC, which we further filtered for genes with prevalence > 0.5% to improve memory footprint and runtime required for profiling) using BWA-MEM (0.7.17) with default parameters and name-sorted by samtools collate (1.14)^45^. Alignments are then filtered to >45bp alignment length and >97% sequence identity, which results in very high mapping rates (**Extended Data Fig. 4**). Cayman quantifies CAZyme domain abundances by counting reads overlapping annotated CAZy domains. Paired-end reads contribute 0.5 counts per mate, reads from single-end libraries contribute 1 count. If reads align to multiple domains, they fractionally contribute towards each domain. To account for biases introduced by gene length and sequencing depth, read counts were normalised to Reads Per Kilobase per Million mapped reads (RPKM) against the number of reads passing the above alignment filters.

### Taxonomic profiling of metagenomic shotgun data

In order to obtain taxonomic profiles, we used mOTUs with default settings (v3.1)^23^. Pre-processing was done as described in the section “Pre-processing of metagenomic datasets”.

### Western and non-Western datasets

To investigate differences in CAZyme repertoire between Western and non-Western populations, we profiled 19 different datasets from 19 different countries and retrieved the definition of Western / non-Western and associated metadata from the curatedMetagenomicData package (v3.6.2)^13^. Binary classification of Western versus non-Western was obtained from this resource and classification is based on the adoption of a Westernised lifestyle encompassing different characteristics, as was previously explained in detail^46^. We exclusively selected samples from healthy controls (n=4,281) also if these were infected with a soil-transmitted helminth (n=108 / 4,281) given the high prevalence of these in many countries in healthy individuals. In cases where individuals were sampled repeatedly, we only retained a single sample (Table S3 for details per dataset).

### Statistical analysis of Western and non-Western data

CAZyme richness was calculated by counting the number of uniquely observed CAZyme (sub)families within each sample, provided that the abundance was > 1 RPKM. Principal coordinates analysis was performed using Canberra distances based on the CAZy RPKM values. Substrate ratios were calculated by summing the RPKMs of all CAZyme families annotated by the given substrate (e.g. DF or GAG) and then computing the ratio between the respective values. CAZyme differential abundance analysis was performed using a linear model implemented in SIAMCAT (v2.5.1)^47^ and obtained P-values were adjusted using the Benjamini-Hochberg method), with adjusted p-values < 0.05 considered significant. Features were filtered based on having a prevalence of at least 1% in the entire dataset prior to performing differential abundance analysis. A pseudocount of 0.01 was added to RPKM values prior to statistical analysis. Generalised fold changes (gFC) were calculated as implemented in SIAMCAT^47^. Considering that almost all studies only had Western or only non-Western samples, we could not block by study.

To study the contribution of microbial taxa to community-level CAZyme abundances in metagenomes, we used the lm function in R to predict log10-scaled CAZyme RPKM values (with a pseudocount of 0.01) based on centred-log ratio scaled species-level microbial profiles of the non-Western/Western dataset collection. For each genus-CAZyme pair, we fit two linear models: One for all non-Western samples and one for all Western samples. For each genus, all mOTUs (i.e. species-level taxonomic groups) of that genus were used to predict CAZyme abundances. All associations computed this way were FDR-corrected using the Benjamini-Hochberg method at 1% FDR and further filtered for those where both models have a coefficient > 0. In the heatmap, we only show those genera with at least one association with R^2^ > 0.4 and those CAZymes with at least one association with R^2^ > 0.4. We finally filtered out all associations where the CAZyme family is genomically absent in the associated genus.

### Colorectal cancer datasets and statistical analysis

We started our meta-analysis by investigating which CAZymes were differentially abundant between CRC and control metagenomes by fitting LMMs with study as a random effect through the SIAMCAT package (**Table S4**). Features with less than 1% prevalence were removed prior to statistical testing. Next, to investigate whether the obtained signatures were consistent across studies, within-study linear models were applied where exclusively the study label (CRC or control) was included as a variable. Generalised fold changes were calculated as implemented in the SIAMCAT package and P-values were adjusted using FDR correction (Benjamini-Hochberg method), with adjusted p-values < 0.05 considered significant. Substrate ratios were calculated in a linear mixed model setting with the Study being included as a random effect. Study-specific ratios can be found in **Extended Data Fig. 10**. In order to compute genera differentially abundant in CRC, we used unpaired, two-sided Wilcoxon tests and FDR-adjusted P-values at 10% using the Benjamini Hochberg method. Given that all studies were case-control studies, no repeated sampling of individuals was performed.

### Gene set enrichment analysis (GSEA)

In order to investigate whether there are substrate preferences among the CAZymes enriched or depleted in CRC metagenomes, we performed GSEA using the R package fgsea (v1.22.0) and the fgseaMultilevel function with default parameters. As input measure for fold change, generalised fold changes as obtained from SIAMCAT were used. Adjusted p-values were calculated to determine significance for each substrate.

### Co-occurrence analysis for CRC-enriched CBMs

In order to compute CAZyme families co-occurring within genes, we computed a modified Jaccard index between all pairs of CAZymes from our CAZy annotations of a large human gut microbial genome collection (see section “Genomic exploration of CAZymes in human gut”). Specifically, we first selected all ORFs that have at least 2 distinct CAZyme families, after which we computed for all pairs of CAZyme families the nominator of the index as the number of ORFs that contain both CAZyme families. For normalisation of this index we computed the cardinality for both CAZyme families separately (in other words, the number of ORFs that contain that given CAZyme family) and normalised the index by the smaller of both values. We computed this index once for each genus containing at least 10 genomes. For the visualisation of this data in Figure 5, we restricted the data to catabolic CAZyme families co-occurring with CRC-enriched CBMs with an index of at least 0.2.

## Data availability

All raw metagenomic data used in this study can be accessed from public repositories through the project numbers listed in the original manuscripts (**Table S3** and **Table S4**). To make our tool as broadly applicable as possible, we annotated all non-human-gut GMGC sub-catalogs excluding genes with less than 0.5% prevalence^18^. All annotated GMGC catalogues are available through Cayman’s GitHub page https://github.com/zellerlab/cayman.

## Code availability

All custom code and required data files to reproduce the analyses and figures can be found at https://git.embl.de/grp-zeller/cazy_gut_microbiome/.

## Acknowledgements

We thank members of the Zeller group for fruitful discussions. We are moreover indebted to the EMBL IT Services HPC for support with high-performance computing. This work received funding from EMBL, the Federal Ministry of Education and Research (BMBF grant no. 031L0181A to G.Z.), the German Research Foundation (Deutsche Forschungsgemeinschaft project number 395357507 – SFB 1371 to G.Z.) and was further supported through a FEMS Research and Training Grant (to Q.D.), the Health + Life Science Alliance Heidelberg Mannheim through state funds approved by the State Parliament of Baden-Württemberg (Postdoctoral Fellowships to Q.D. and N.K.) and an EMBO postdoctoral fellowship (EMBO ALTF 1030-2022 to Q.D.)

## Authors’ contributions

QD, NK and GZ conceived and designed the project. QD and NK performed all bioinformatic and statistical analyses. QD, NK and GZ designed figures and drafted the manuscript. QD, HT and ODB performed substrate annotations. FS and SP performed taxonomic profiling of metagenomic cohorts and aided in curating metadata. CS designed and engineered the Cayman software. GZ supervised all aspects of this work.

## Supple

**Supplementary Table 1:** Hierarchical substrate annotations for all CAZyme families of the classes GH, PL and CBM.

**Supplementary Table 2:** Table with glycans and their hierarchical substrate annotations.

**Supplementary Table 3**: List of the studies included in our Western / non-Western meta-analysis and notes associated with metadata curation for each of the studies.

**Supplementary Table 4**: List of studies included in our CRC meta-analysis.

