## Extended_Data_Figures for "Large-scale computational analyses of gut microbial CAZyme repertoires enabled by Cayman"

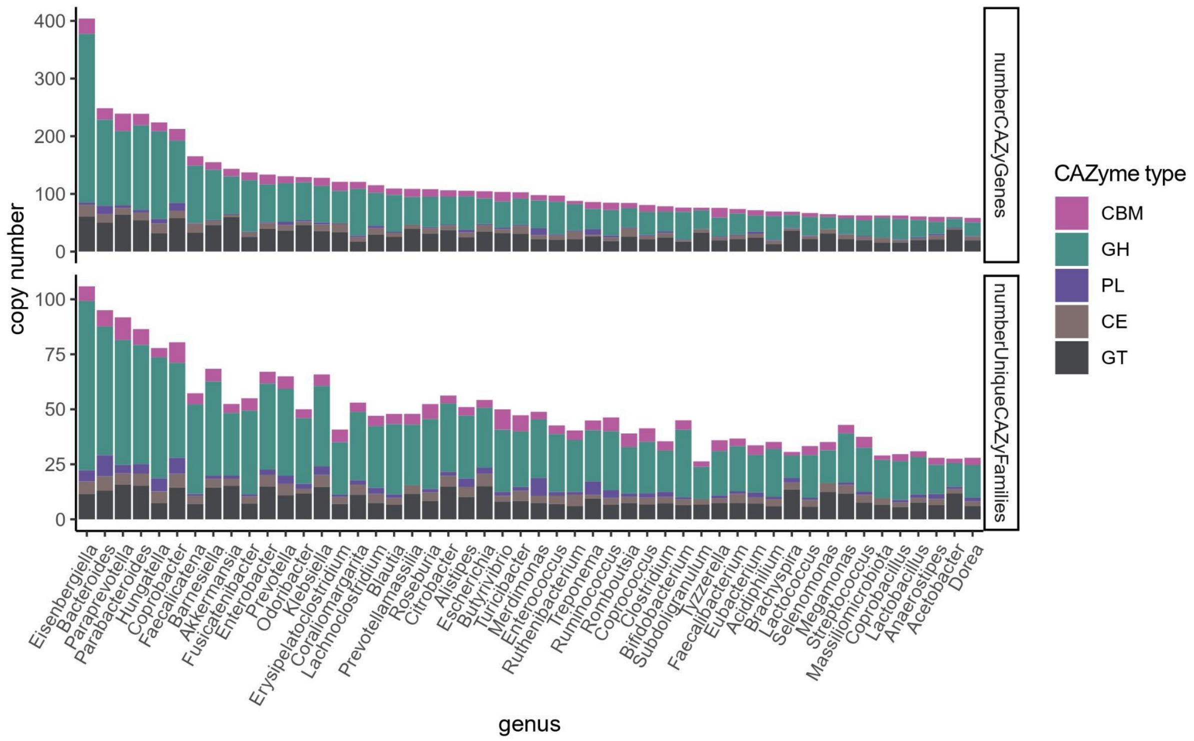


**Extended Data Fig. 1: Copy number of CAZymes in bacterial genera.** Total number of CAZyme genes and number of unique CAZyme families per genus. GH; Glycoside Hydrolase, PL; Polysaccharide Lyase, CE; Carbohydrate Esterases, GT; Glycosyl transferases and CBM; Carbohydrate Binding Module.

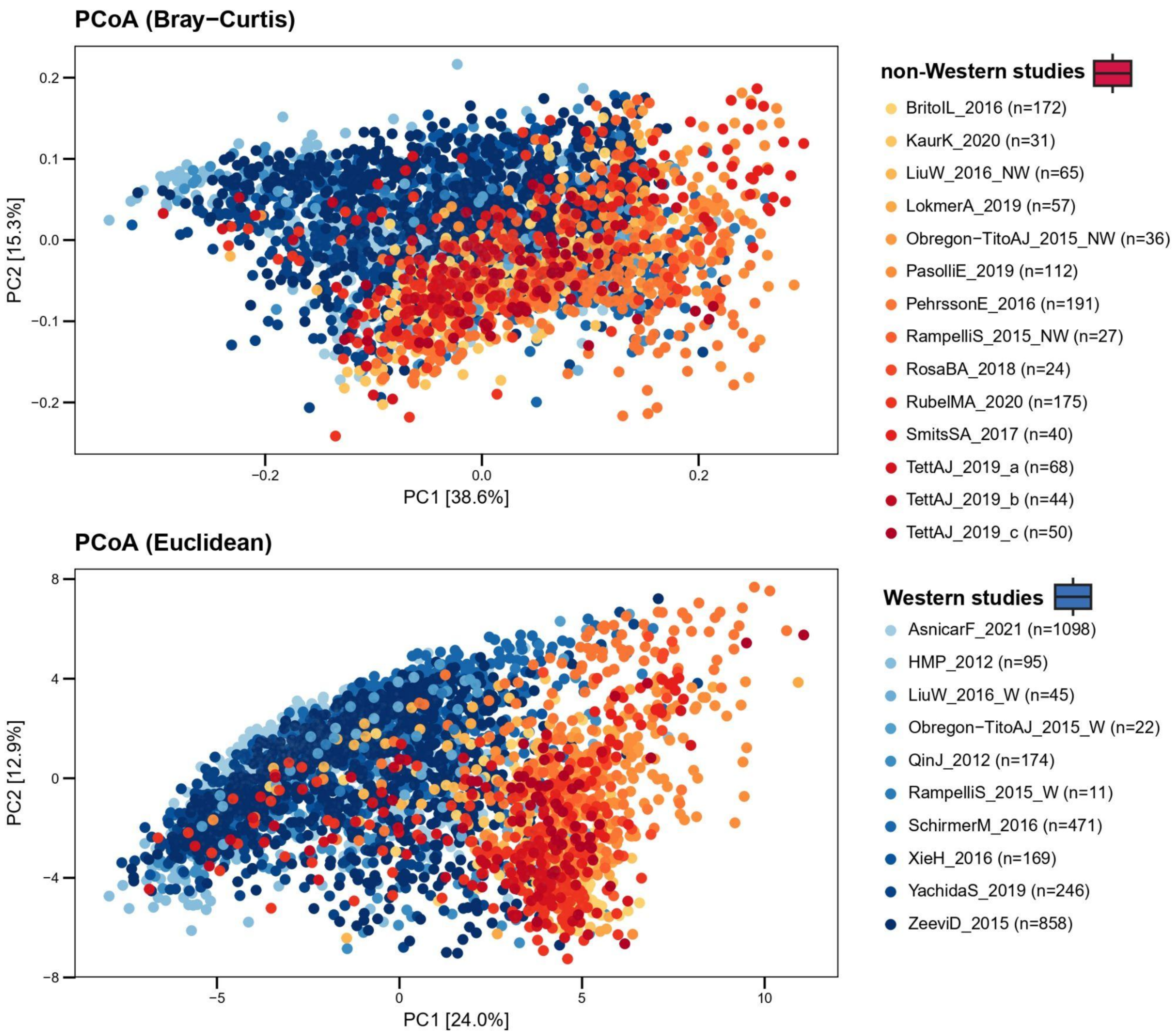


**Extended Data Fig. 2**: **Principal Coordinates Analysis**. Principal Coordinates Analysis of 4,281 Western- and non-Western metagenomes using Bray-Curtis dissimilarity and log-transformed Euclidean distances. Corresponds to Fig. 3A.

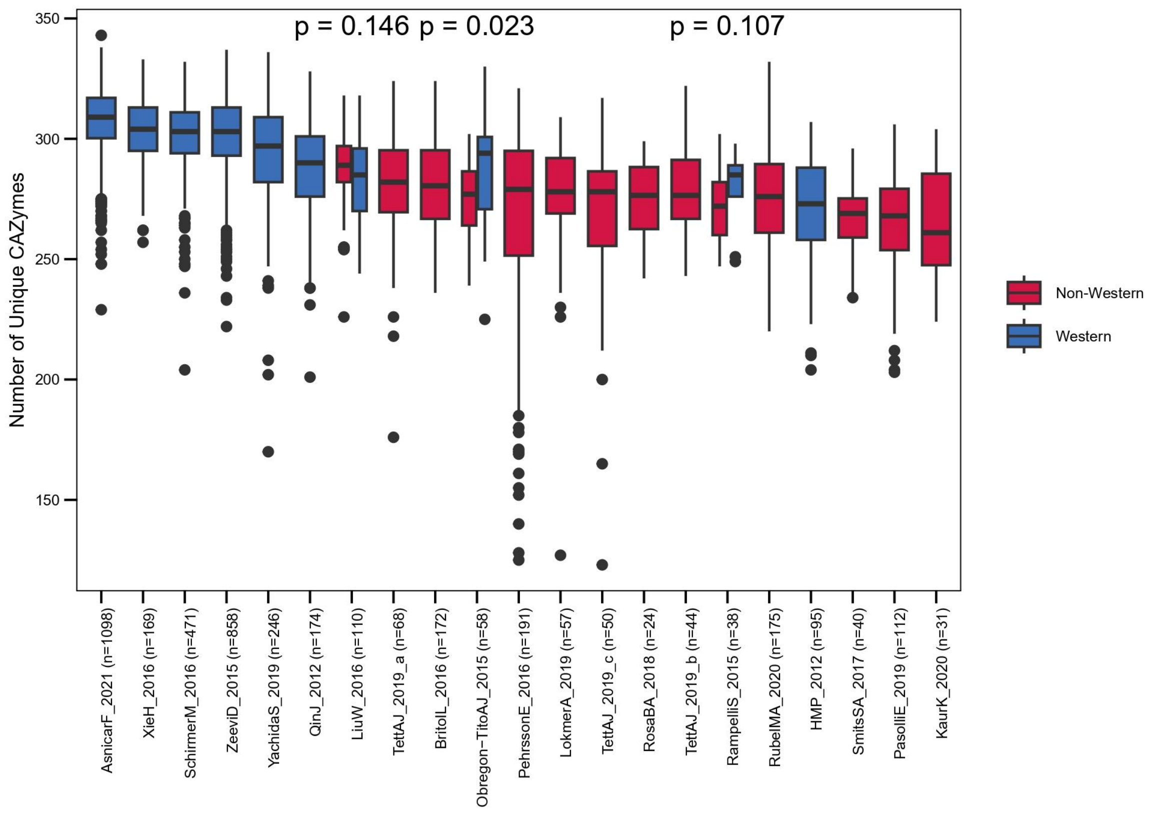


**Extended Data Fig. 3: Number of unique CAZyme families across all studies.** CAZyme richness for all studies included in our Western versus non-Western meta-analysis. Studies are ordered by the median CAZyme richness. P-values were computed using unpaired, two-sided Wilcoxon tests for the studies that had both Western and non-Western samples included.

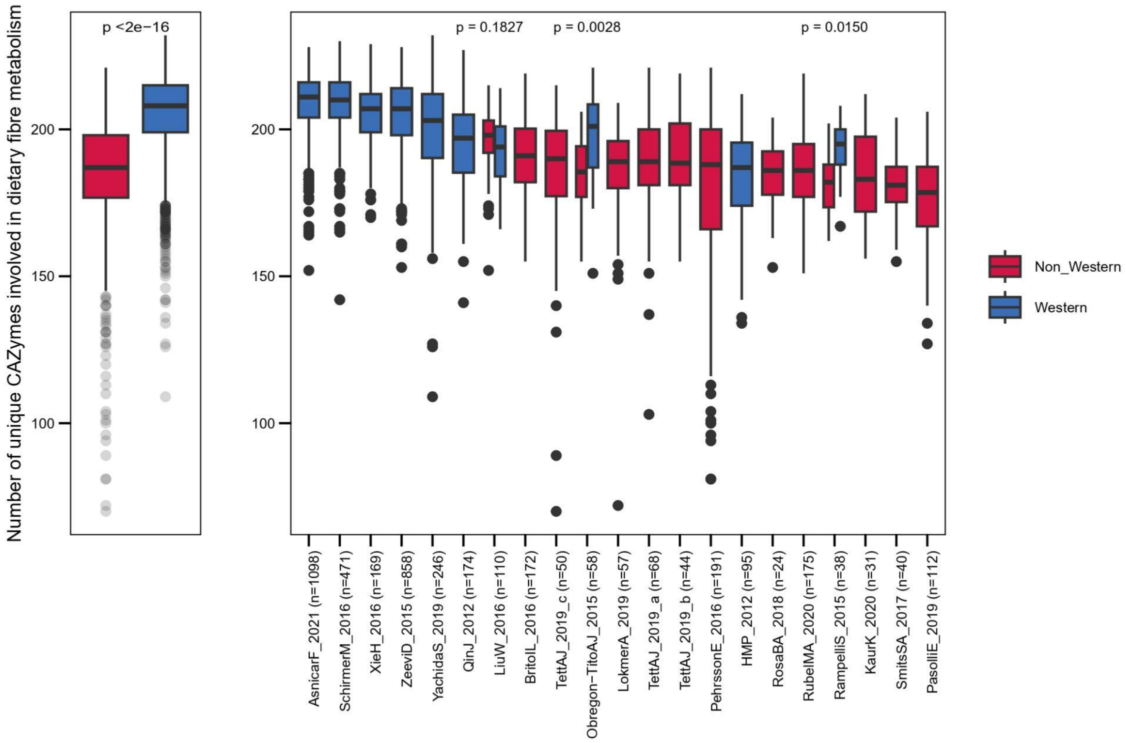


**Extended Data Fig. 4: Number of unique dietary-fibre targeting CAZyme families across all studies.**

CAZyme richness of only CAZymes involved in dietary fibre metabolism for all studies included in our Western versus non-Western meta-analysis, as well as the two groups combined. Studies are ordered by the median CAZyme richness. P-values were computed using unpaired, two-sided Wilcoxon tests for the studies that had both Western and non-Western samples included.

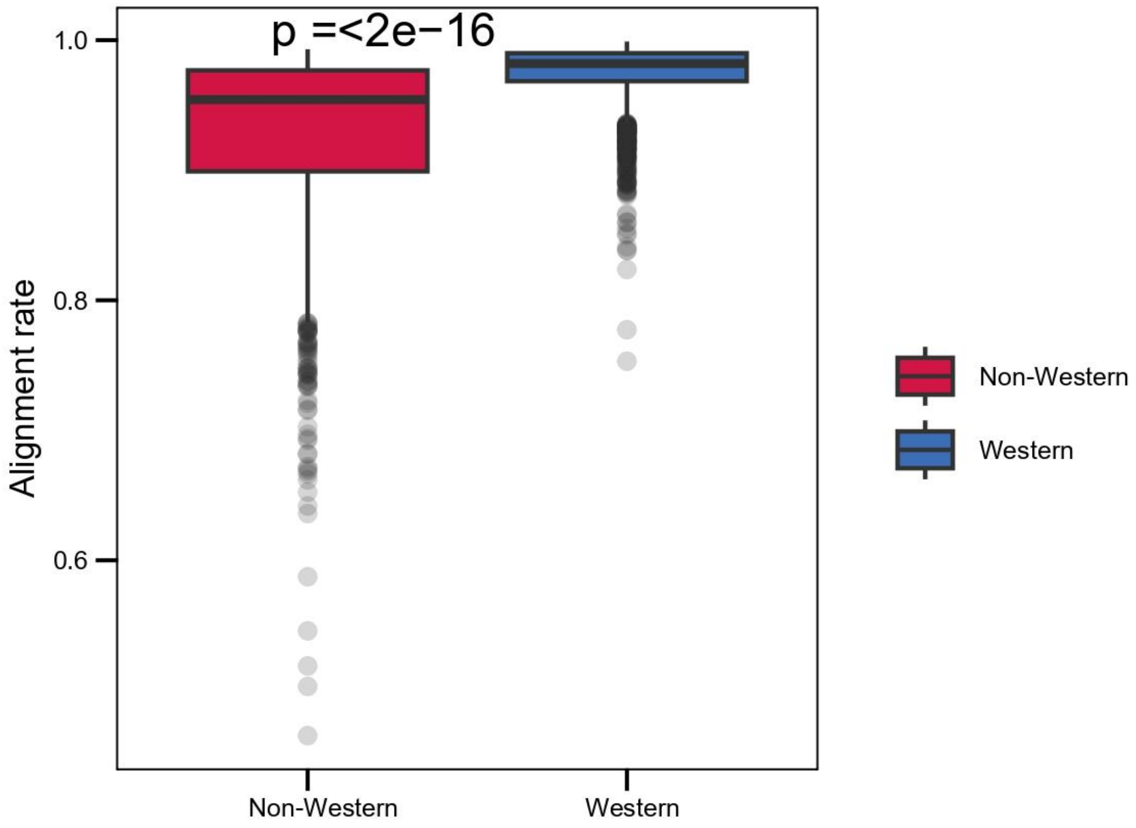


**Extended Data Fig. 5: Alignment rates to our gut GMGC catalogue.** Boxplot of metagenome alignment rates against the GMGC catalogue used for CAZyme profiling. P-value was computed using an unpaired, two-sided Wilcoxon test.

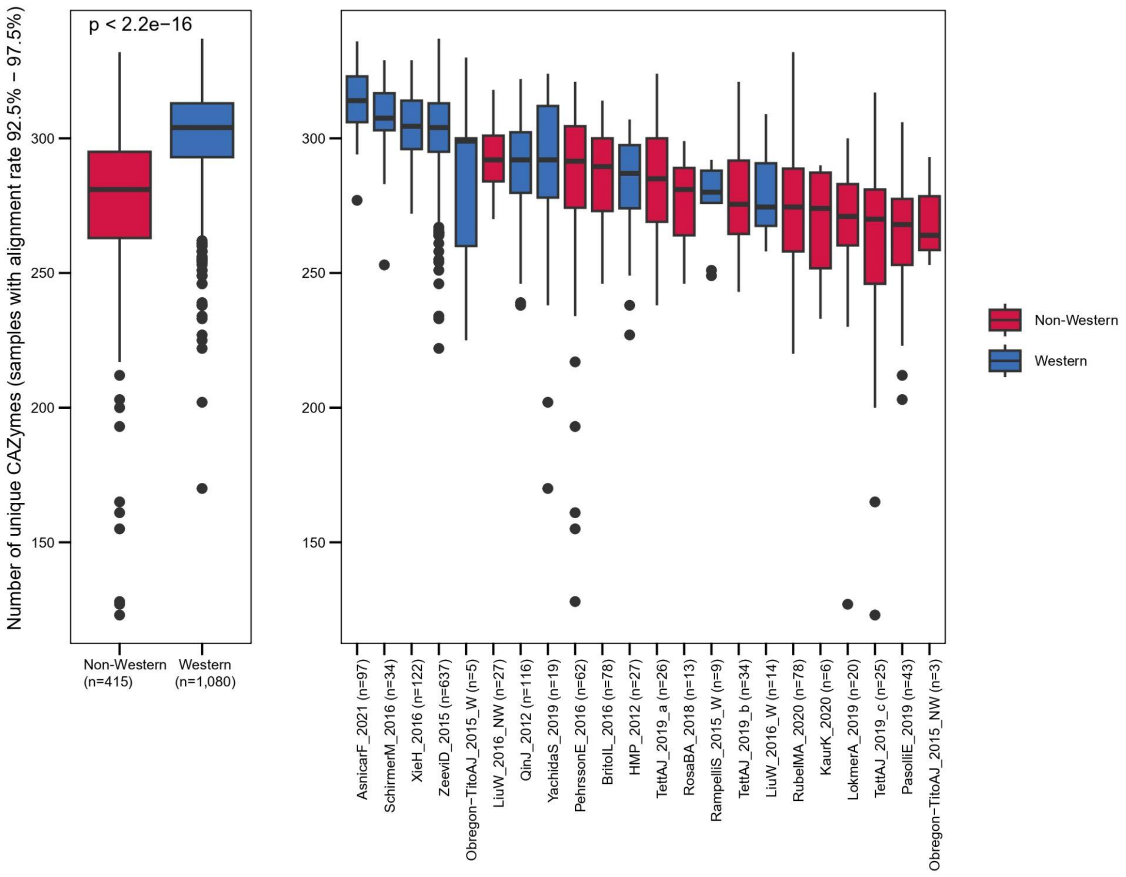


**Extended Data Fig. 6: Number of unique CAZyme families on samples with alignment rates between 92.5% and 97.5%.** CAZyme richness in Western/non-Western metagenomes with high alignment rates (92.5% - 97.5%). Studies are ordered by the median CAZyme richness. P-value was obtained using an unpaired, two-sided Wilcoxon test.

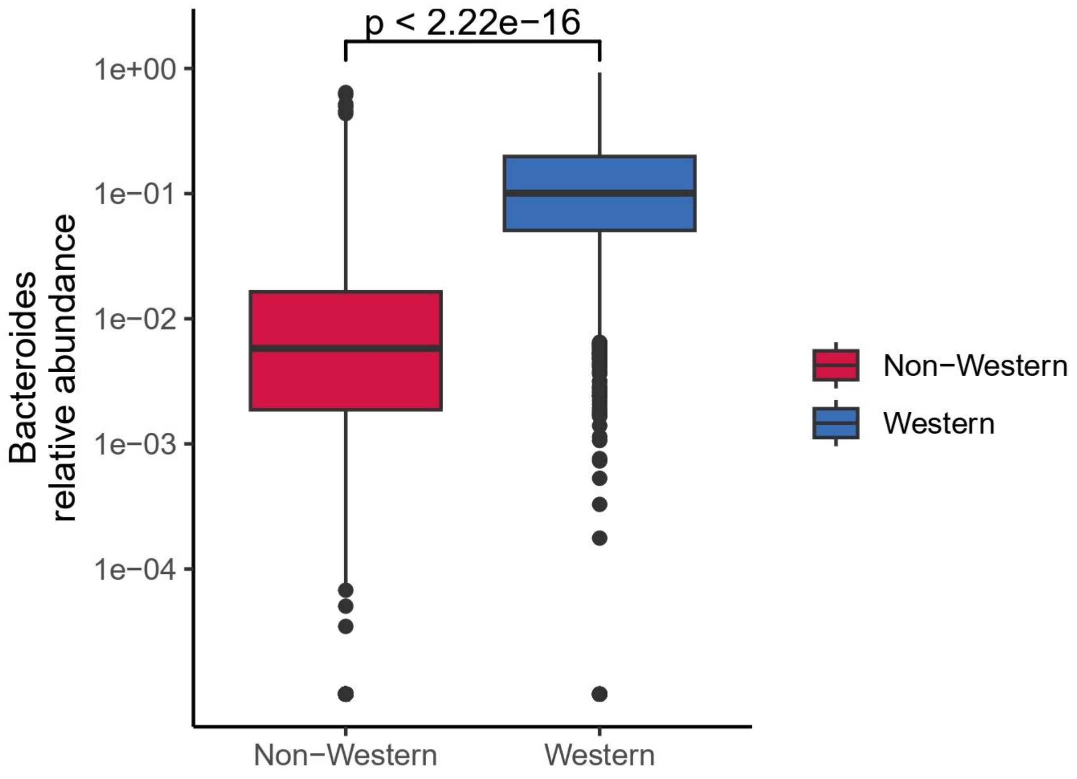


**Extended Data Fig. 7: *Bacteroides* relative abundance.** Relative abundance of *Bacteroides* in Western versus non-Western individuals. P-value was obtained using an unpaired, two-sided Wilcoxon test.

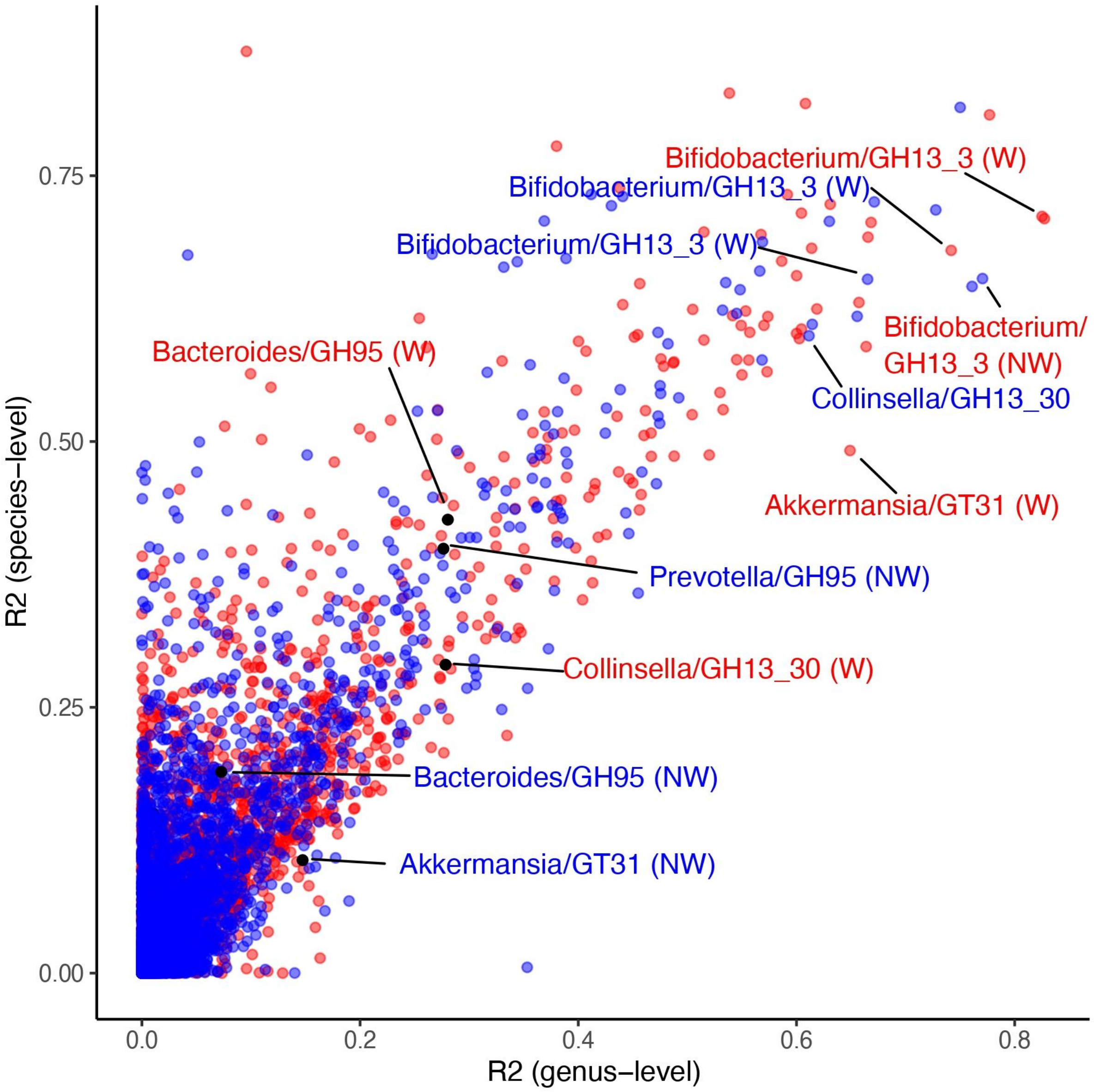


**Extended Data Fig. 8: Scatter plot of linear model R2 values predicting CAZyme family abundances from genus- and species-level taxonomic profiles.** X-axis corresponds to models using genus-level microbial abundance as predictor; Y-axis corresponds to models using mOTU-level microbial abundances as predictor. Label family/genus combinations correspond to examples illustrated in Fig. 4B-4G.

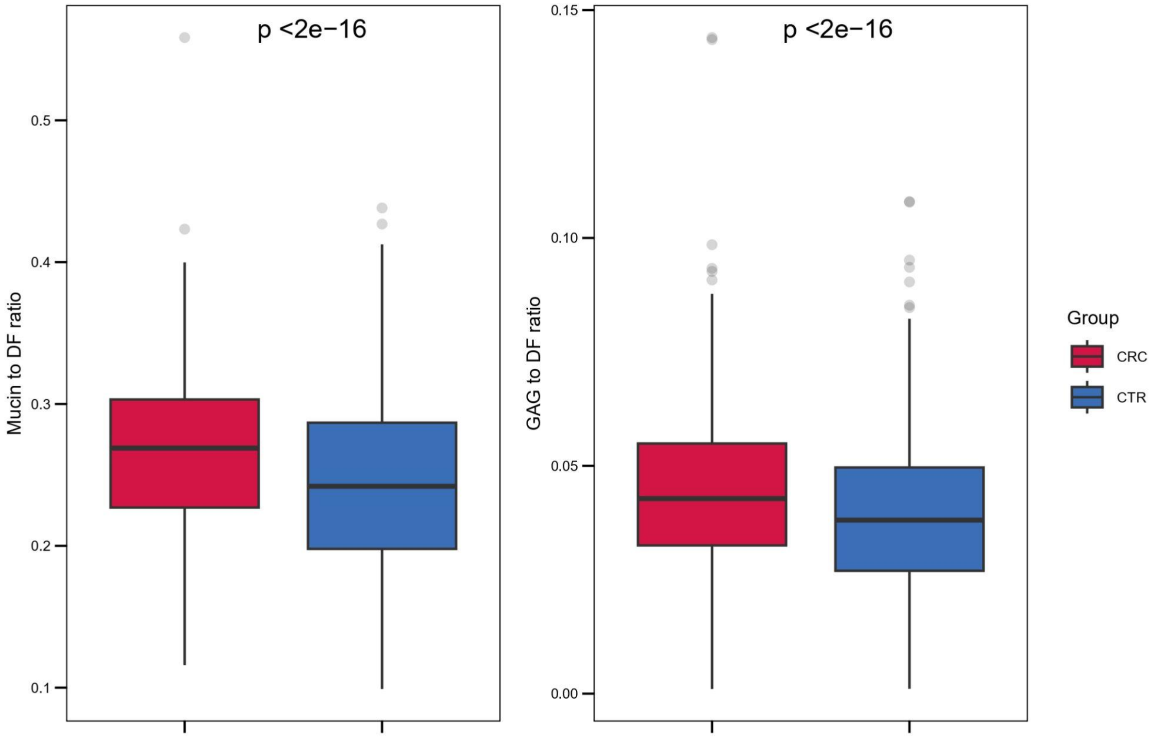


**Extended Data Fig. 9: CAZyme Substrate ratios in colorectal cancer patients versus controls.** Boxplots of the total abundance of mucin-targeting CAZymes and GAG-targeting CAZymes, each normalised by the total abundance of dietary fibre-targeting CAZymes in CRC patients versus controls. P-values were obtained using unpaired, two-sided independent Wilcoxon tests. CRC; Colorectal cancer, CTR; Controls.

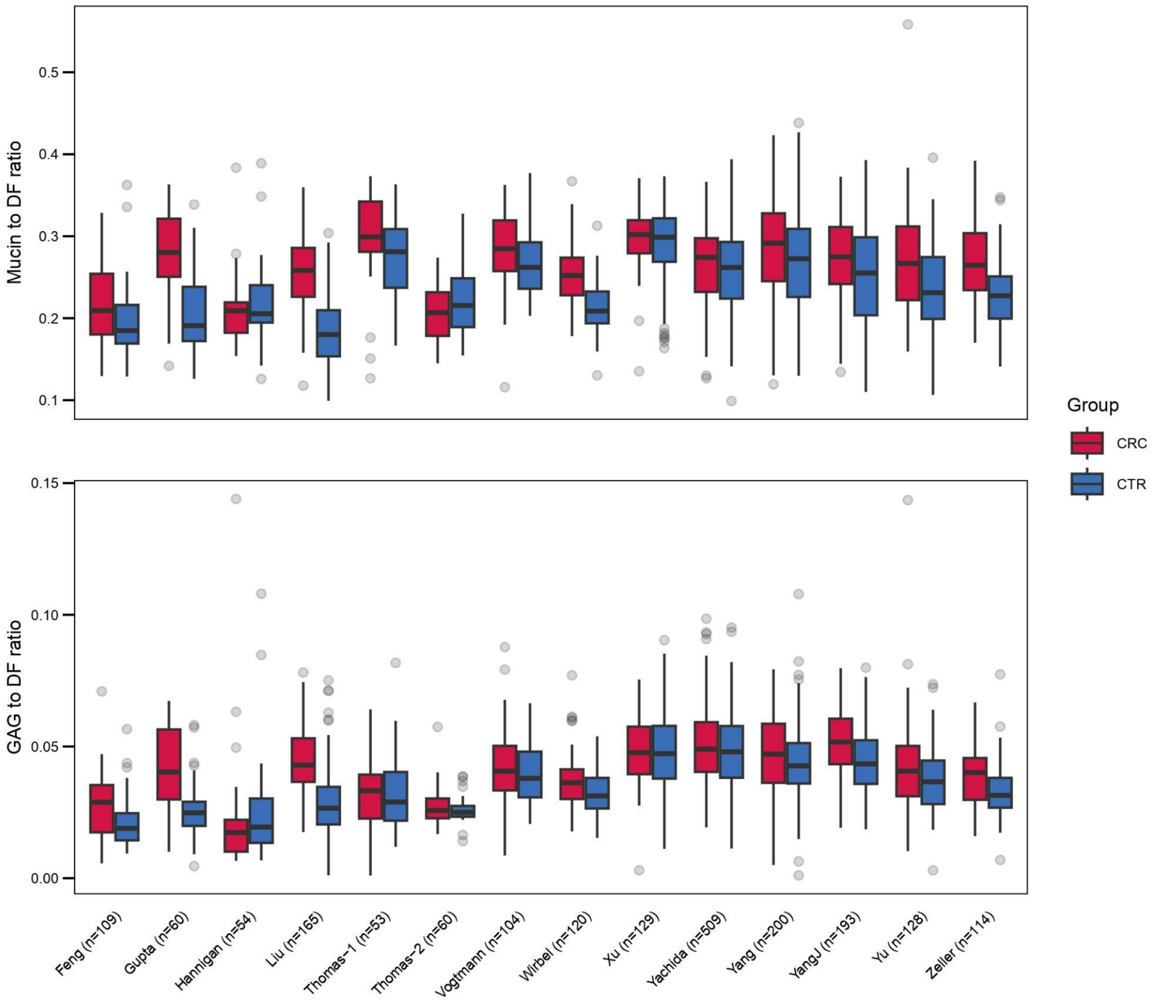


**Extended Data Fig. 10: CAZyme substrate ratios in colorectal cancer patients versus controls for all studies.** Boxplots of the total abundance of mucin-targeting CAZymes and GAG-targeting CAZymes, each normalised by the total abundance of dietary fibre-targeting CAZymes in CRC patients versus controls. CRC; Colorectal cancer, CTR; Controls.
